# Boosting L-type Ca^2+^ channel activity in tuberous sclerosis mitigates excess glutamate receptors

**DOI:** 10.1101/2020.08.26.269209

**Authors:** Farr Niere, Luisa P. Cacheaux, Hailey X. Egido-Betancourt, William C. Taylor, Kimberly F. Raab-Graham

## Abstract

Tuberous sclerosis complex (TS) is a dominant, multisystem disorder with devastating neurological symptoms. Approximately 85% of TS patients suffer from epilepsy over their lifespan and roughly 25-50% of those patients develop Autism Spectrum Disorder (1, 2). Current seizure therapies are effective in some, but not all, and often have significant risk factors associated with their use (1, 3). Thus, there is a critical need for new medication development or drug repositioning. Herein, we leveraged proteomic signatures of epilepsy and ASD, often comorbid in TS, to utilize an *in silco* approach to identify new drug therapies for TS-related seizures. We have discovered that activation of L-type voltage dependent calcium channels (L-VDCC) by Bay-K8644 (BayK) in a preclinical mouse model of TS rescues the excess expression of ionotropic, AMPA-subtype glutamate (GluA) receptors. As added proof of BayK working through L-VDCC to regulate GluA levels, we found that increasing expression of alpha2delta2 (α2δ2), an auxiliary calcium channel subunit that boosts L-VDCC surface expression, similarly lowers the surface expression of dendritic GluA in TS. These BayK-induced molecular alterations may potentially improve the quality of life of humans suffering from TS.

**Significance Statement:** Causal mechanisms of Tuberous Sclerosis (TS)-associated neurological disorders are under-characterized and treatment options are lacking. Using a computational approach of mTOR/DJ-1 target mRNAs to predict new medications, we report that boosting L-type voltage-dependent Ca^2+^ channel (L-VDCC) activity in a preclinical TS mouse model that exhibits a deficit in dendritic L-VDCC activity ameliorates key molecular pathologies that are predicted to underlie seizures. Restoring the mTOR/DJ-1 pathway upstream of L-VDCC, therefore, may serve as a new therapeutic avenue to mitigate seizures and mortality in TS.

## Introduction

The majority of patients with TS struggle with neurological and neuropsychiatric problems throughout their lifetime (4). The etiology of TS stems from loss-of-function mutations of either the *TSC1* or *TSC2* gene. Normally, Tsc1 and Tsc2 proteins work in conjunction to limit the activity of the mammalian target of rapamycin complex 1 (mTORC1), a key signaling pathway in protein homeostasis (5). Numerous investigations on mTORC1 demonstrate that its activity can equally promote and inhibit protein expression, especially in synaptic regions (6-10). Emerging studies now also indicate that persistent over- or underactive mTORC1 state can steer neuronal and network excitability towards pathological conditions, as mTORC1 regulates the expression of ion channels and their associated proteins (7, 11-13) *(see accompanying manuscript)*. Indeed, several diseases—epilepsy and autism spectrum disorder (ASD) to name a few—that exhibit persistent mTORC1 activity also have enduring states of neuronal or network hyperexcitability (14-17). A mechanism by which mTORC1 can influence ion channel expression and activity, and potentially behavior, is by controlling the levels of RNA-binding proteins (RBPs). RBPs profoundly impact the translation and/or stability of target mRNAs (5, 9, 18) *(see accompanying manuscript)*. By regulating the expression or repression of their target mRNAs, RBPs have the capacity to coordinate the synaptic function (5, 19). We have recently discovered that mTORC1 can dampen dendritic L-type voltage-dependent calcium (Ca^2+^) channel (L-VDCC) activity through the RNA-binding protein DJ-1 (encoded by *Park7*) *(see accompanying manuscript)*. In a preclinical, mouse model of TS, in which *Tsc1* has been conditionally knocked-out (*Tsc1* KO), DJ-1 expression is elevated in dendrites and postsynaptic density (PSD)-positive regions (9). In light of the changes in DJ-1’s subcellular localization in response to mTORC1 activity and DJ-1’s ability to control protein expression of its target mRNAs, we explored the therapeutic potential of mitigating DJ-1/L-VDCC signaling in TS.

## Results

### Prediction of Bay-K-8644 to oppose the effects of aberrant mTOR/DJ-1 at the synapse

As a first step in evaluating if DJ-1 plays a relevant role in TS and the PSD, we searched the mRNA transcripts that encode PSD-associated proteins for DJ-1 binding sites (9) *(see accompanying manuscript)* (**Table S1**). Remarkably, ∼13% of risk genes for TS-associated neurological disorders (epilepsy and autism spectrum disorder (ASD)) are PSD-associated proteins that are encoded by mRNAs possessing DJ-1-binding motifs (**Fig. 1A; Tables S2 and S3**). With putative DJ-1-target mRNAs in hand, we used an unbiased, bioinformatics approach to pinpoint specific DJ-1 function and to identify potential new therapeutic reagents. To accomplish this, we used connectivity mapping (CMap)—an algorithm that can be used to predict drugs that may match or oppose a given change in a gene-expression profile (20). We hypothesized that DJ-1-target mRNAs—predicted to be repressed when mTORC1 is overactive—are homeostatic and neuroprotective. Inputting putative DJ-1-target mRNAs that are predicted to go up when mTOR is off, surprisingly identified Bay-K-8644 (BayK)—an L-VDCC opener—as a candidate-drug with a score of 97 (**Table S4**). A tau score that is higher than 90 is generally considered a compelling candidate for further study (21). Also in line with our hypothesis, L-VDCC is thought to mediate homeostasis (22).

**Figure 1.**
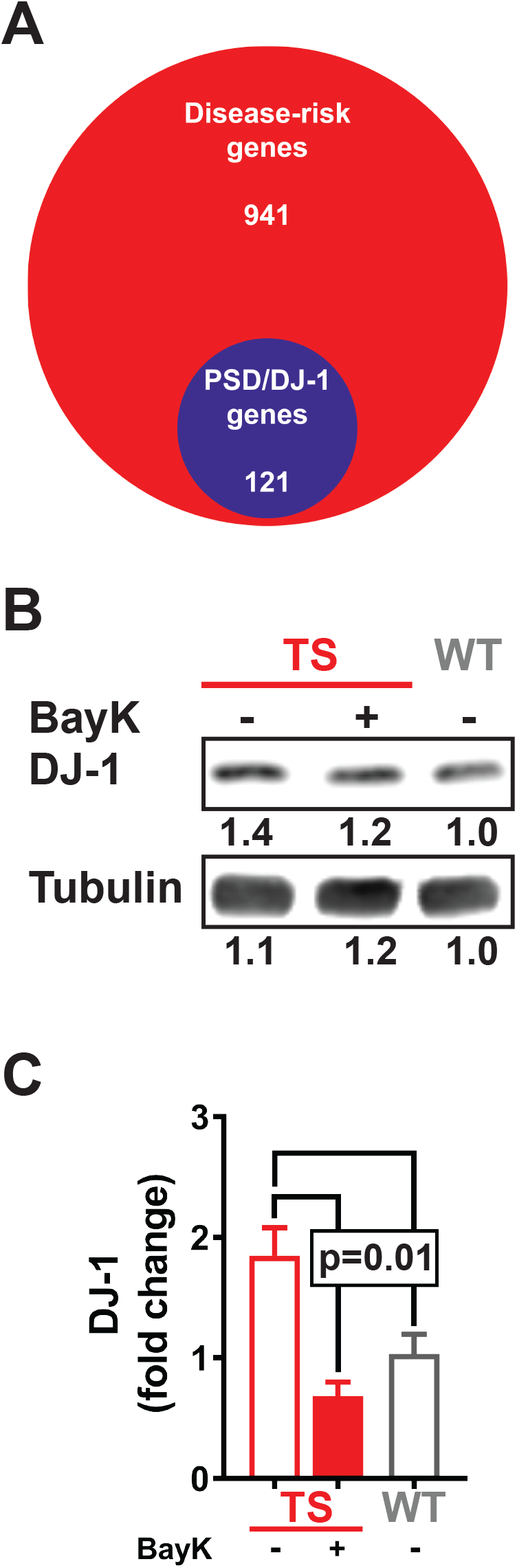
Boosting L-type VDCC activity by Bay-K-8644 (BayK) in a mouse model of tuberous sclerosis (TS) restores excessive DJ-1 expression. **(A)** 121 or 13% of epilepsy- and ASD-risk genes are associated with the PSD and contain several putative DJ-1-binding sequences (GNGCNG and CNGCNG). **(B and C)** Western blot experiments demonstrate that 18-20 hours post-administration of BayK (0.2 to 2 mg/kg) in TS mice lowers synaptic DJ-1 to wildtype (WT) level. **(B)** Representative Western blot of DJ-1 and tubulin and their normalized densitometric values provided below each band. Values are normalized to WT. **(C)** Quantification of Western blot experiments normalized to WT (TS: 1.85±0.23, n=4; TS+BayK: 0.68±0.12, n=4; WT: 1.00±0.16, n=7; one-way ANOVA). Values are shown as mean±s.e.m.

CMap analysis suggests that treating TS with BayK will reverse DJ-1 expression—returning it back to WT levels—and restore L-VDCC-mediated functions despite the significant deficit in dendritic L-type activity *(see accompanying manuscript)*. To test this prediction, we generated a preclinical mouse model of TS, whereby *Tsc1* was conditionally knocked out (*Tsc1* KO) in *Tsc1*^*tm1Djk*^*/J* mice by stereotaxically introducing AAV-Cre-GFP or AAV-GFP, which serves as control, in the hippocampi. mTORC1 activity and DJ-1 levels in dendrites and PSD-positive regions are elevated in TS mice—those that received Cre-GFP—compared to controls (9). We then gave TS mice a single injection of BayK (0.2-2mg/kg) intraperitoneally (i.p.) and measured synaptic DJ-1 protein levels 18-20 hours post-injection. Results of western blot assays indeed indicate that BayK treatment can restore DJ-1 protein levels in TS synaptoneurosomes down to WT level, as predicted by CMap analyses (**Fig. 1B**).

### BayK restores elevated membrane-surface GluA in TS to wildtype condition

L-VDCC performs many cellular functions, including promoting endocytosis of AMPA-subtype glutamate receptors (GluA) (23). It’s been reported that GluA1 levels are elevated in the human condition of TS (24). Moreover, GluA-mediated currents are enhanced in a *Tsc1* KO mouse model (25, 26). To mimic L-VDCC hypofunction in TS and to assess the impact of reduced L-type channel activity on GluA expression, we blocked L-type activity of dissociated WT hippocampal neurons with 5 μM nimodipine for 30 minutes and subsequently measured the surface expression of GluA (sGluA) on dendrites by immunocytochemistry. Of note, sGluA levels correlate well with GluA-mediated currents (27-29). L-VDCC blockade by nimodipine increases sGluA by ∼twofold over control (**Fig. 2A and B**). Building on this result, we treated dissociated hippocampal neurons from TS mice with 5 μM BayK for 60 minutes and assessed sGluA expression. Our results show that TS neurons, compared to WT, express twofold more sGluA. Furthermore, applying BayK to TS neurons lowers their sGluA levels to WT state (**Fig. 2C and D**).

**Figure 2.**
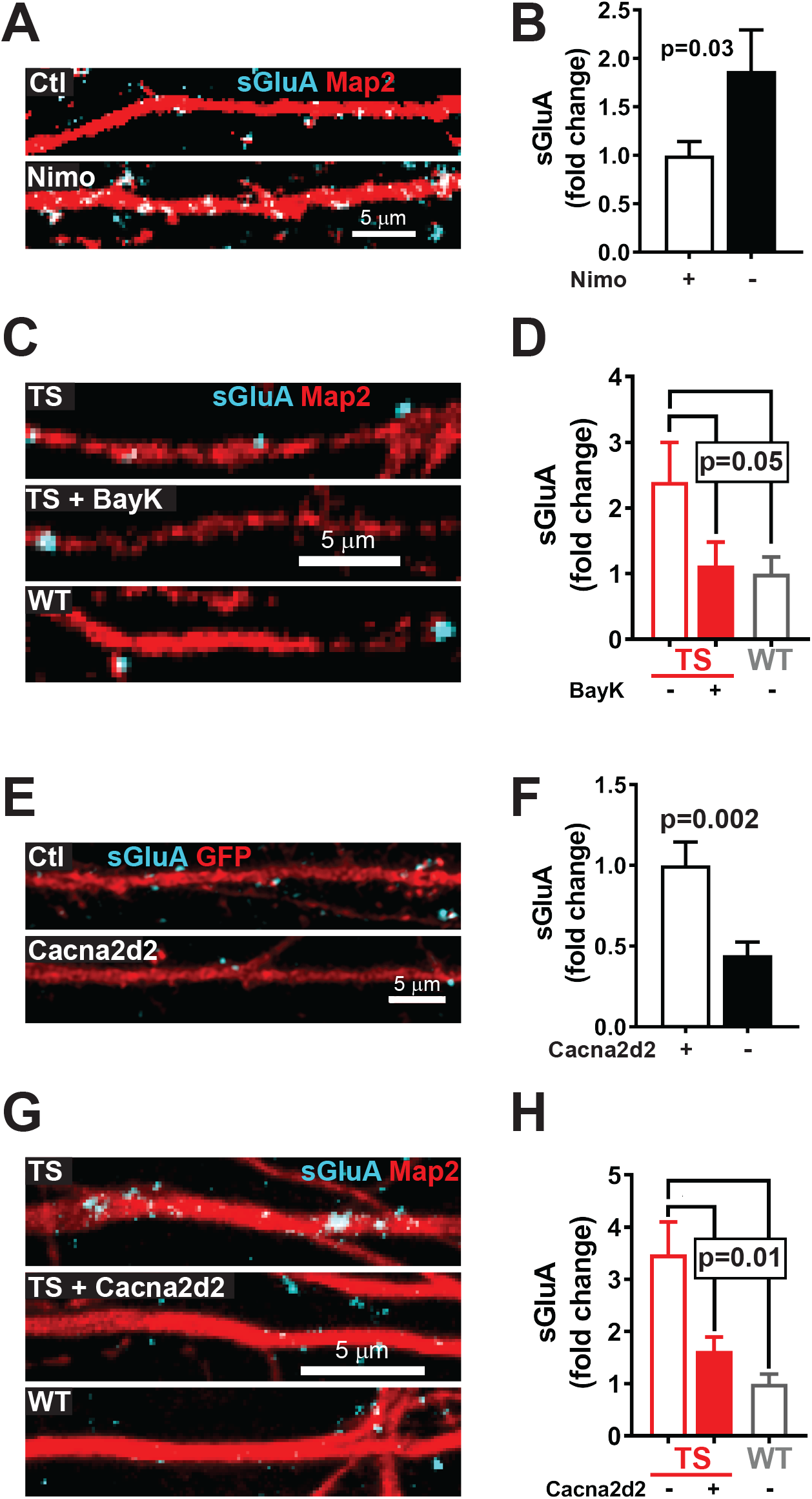
BayK or overexpression of calcium channel-associated protein α2δ2 in TS neurons corrects the excessive expression of GluA on the dendritic membrane. **(A-D)** L-type calcium channels control the surface expression of GluA (sGluA). **(A-B)** Dissociated rat hippocampal neurons treated with L-type VDCC blocker nimodipine (Nimo; 5 μM) have fewer sGluA compared to control (Ctl) as shown by **(A)** representative figures and **(B)** quantification of sGluA normalized to Ctl (Ctl: 1.00±0.14, n=19; Nimo: 1.87±0.42, n=13; student’s t-test). **(C-D)** Excess sGluA levels in TS can be curtailed by application of BayK (5 mM) to WT status as depicted in **(C)** representative figures and **(D)** quantification of sGluA normalized to WT (TS: 2.54±0.60, n=18; TS+BayK: 1.12±0.36, n=20; WT: 1.00±0.26, n=16; one-way ANOVA). **(E-H)** Levels of *Cacna2d2* gene, which encodes α2δ2 protein, regulate sGluA expression. **(E-F)** Overexpressing *Cacna2d2* in dissociated rat hippocampal neurons diminishes sGluA levels compared to Ctl as shown by **(E)** representative figures and **(F)** quantification of sGluA normalized to Ctl (Ctl: 1.00±0.15, n=25; *Cacna2d2*: 0.44±0.08, n=25; student’s t-test). **(G-H)** Overexpression of *Cacna2d2* in TS lowers elevated sGluA to WT level as depicted in **(G)** representative figures and **(H)** quantification of sGluA normalized to WT (TS: 3.48±0.62, n=14; TS+*Cacna2d2*: 1.63±0.26, n=19; WT: 1.00±0.19, n=12; one-way ANOVA). Map2 indicates dendrites. Values are shown as mean±s.e.m.

### Overexpression of DJ-1-repressed α2δ2 restores elevated membrane-surface GluA in TS to WT condition

Since DJ-1 lowers the dendritic level of α2δ2, which facilitates the surface membrane expression of pore forming Ca^2+^ channel subunits and increases current density, we investigated the influence of α2δ2 on sGluA expression (30-32). TS dendrites notably have more GluA but less α2δ2 than WT *(see accompanying manuscript)*. We therefore considered the possibility that overexpressing α2δ2 in dissociated rat neurons would curtail sGluA expression. Overexpressing *Cacna2d2*— the gene that encodes α2δ2 protein—as predicted lowers sGluA expression (**Fig. 2E and F**). Additionally, increasing α2δ2 levels in TS restores sGluA to WT level (**Fig. 2G and H**). These findings collectively demonstrate that DJ-1 can modulate postsynaptic function through L-VDCC.

### In vivo administration of BayK ameliorates the exaggerated protein expression of GluA and ribosomal S6 in TS

As our *in vitro* data indicate that BayK can mitigate the overabundance of GluA in TS, we assessed BayK’s effect *in vivo*. TS mice were treated with vehicle or the same concentration of BayK that lowers DJ-1 expression to WT level (**Fig. 1B**). Eighteen to twenty hours post-injection, we measured a population of GluA that is phosphorylated at serine 845 residue (pGluA1), which is associated with increased GluA-mediated synaptic current (33). By western blot assays of hippocampal synaptoneurosomes, pGluA1 expression in BayK-treated mice showed a direction towards WT condition, although it did not reach significance (**Fig. 3A and B**). Total GluA expression, however, showed a marked reduction in BayK-injected TS mice compared to vehicle-treated (**Fig. 3C**). Because increases in GluA expression is sensitive to mTOR activity, we examined whether BayK can dampen mTOR activity by measuring the levels of phospho-S6 (pS6)—a downstream effector of mTOR activity—in vehicle and BayK-treated TS animals (34, 35). BayK does reduce pS6 expression in TS; but, BayK does not lower pS6 to WT levels (**Fig. 3D and E**). BayK, however, curtails total S6 to WT condition (**Fig. 3F**). These data suggest that BayK possesses a therapeutic value in TS, as a single peripheral administration of BayK can correct GluA expression by partially repressing mTORC1 activity.

**Figure 3.**
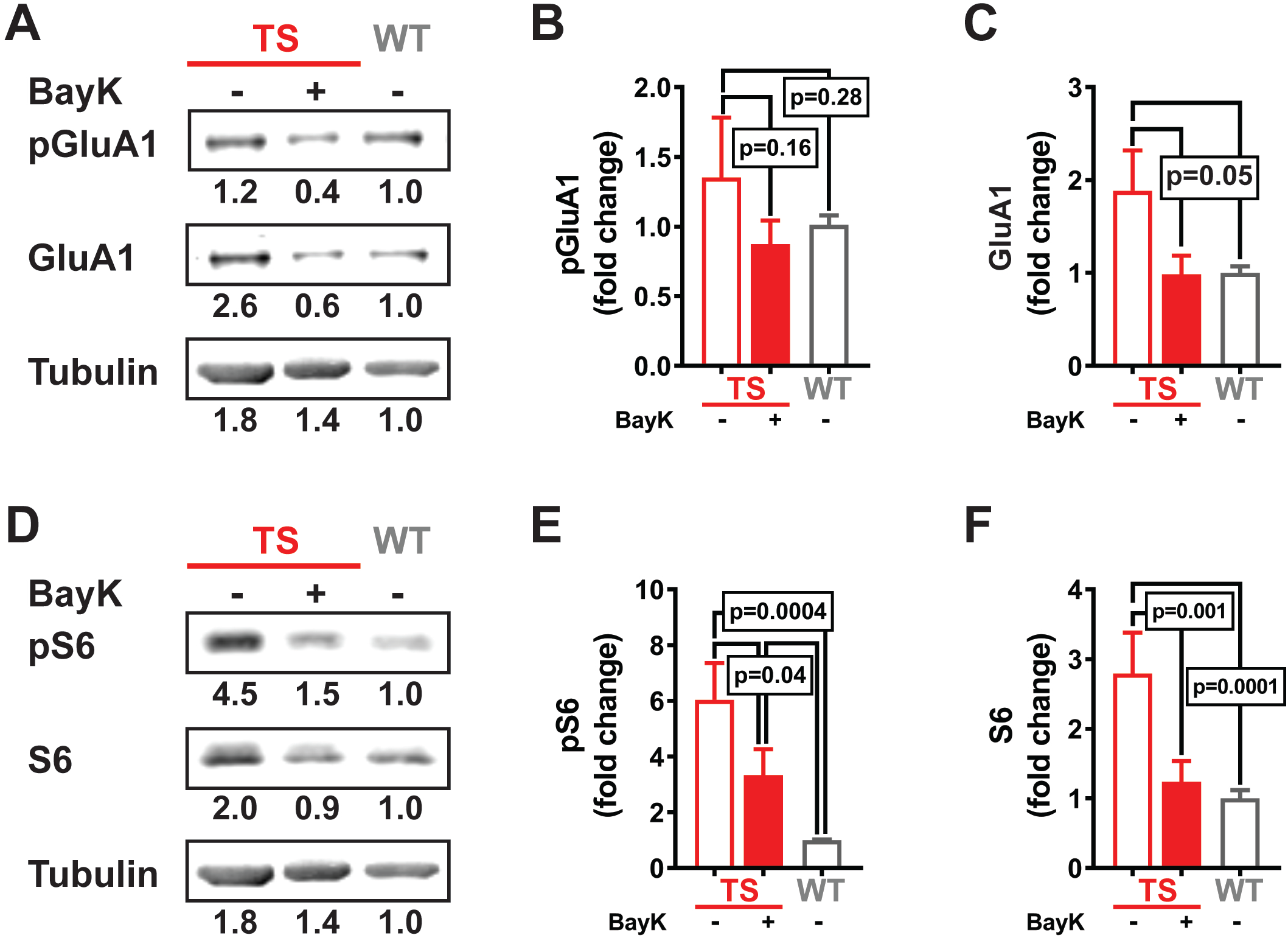
Peripheral administration of BayK in TS mitigates key components of seizure neuropathophysiology. **(A-C)** Acute intraperitoneal (i.p.) injection of BayK (0.2-2 mg/kg) lowers synaptic GluA in TS mice to WT status. **(A)** Representative Western blot of GluA1 phosphorylated at serine 845 (pGluA1), total GluA1 and tubulin and their normalized densitometric values indicated below each band. Values are normalized to WT. **(B)** Quantification of pGluA1 (TS: 1.35±0.43, n=4; TS+BayK: 0.87±0.38, n=5; WT: 1.00±0.18, n=7; one-way ANOVA). **(C)** Quantification of GluA1 (TS: 1.89±0.43; TS+BayK: 0.99±0.45; WT: 1.00±0.19). **(D-F)** Acute i.p. injection of BayK (0.2-2 mg/kg) lowers synaptic ribosomal S6 (S6) protein in TS to WT status. **(D)** Representative Western blot of phosphorylated S6 (pS6), total S6 and tubulin and their normalized densitometric values indicated below each band. Values are normalized to WT. **(E)** Quantification of pS6 (TS: 6.04±1.32, n=4; TS+BayK: 3.34±0.93, n=5; WT: 1.00±0.02, n=7; one-way ANOVA). **(F)** Quantification of S6 (TS: 2.79±0.59; TS+BayK: 1.24±0.30; WT: 1.00±0.12). Values are shown as mean±s.e.m.

## Discussion

The symptoms of TS include epilepsy, ASD and intellectual disability (1, 36-38). As these symptoms implicate postsynaptic dysfunctions, our PSD-targeted, bioinformatics approach using CMap reveals that activating L-type VDCC may mitigate TS-associated neurological disorders (**Fig. 1**). This finding is indeed surprising as L-type channel agonists are rarely considered as therapeutic. Administered to normal organisms, L-VDCC activation can induce neurological and neuropsychiatric symptoms (39). In TS where L-type channel activity is low at best, agonism of the channel is beneficial as it can reduce excess GluA receptors from the membrane surface (**Fig. 2**). The data we present here posits a working model that accounts the molecular changes occurring postsynaptically when mTOR switches from being dynamic (**Fig. 4A**) to being persistently active (**Fig. 4B**). While an overactive mTOR state greatly impedes L-VDCC activity, boosting L-type channels can ameliorate the molecular dysfunctions that may underlie TS-associated seizures.

**Figure 4.**
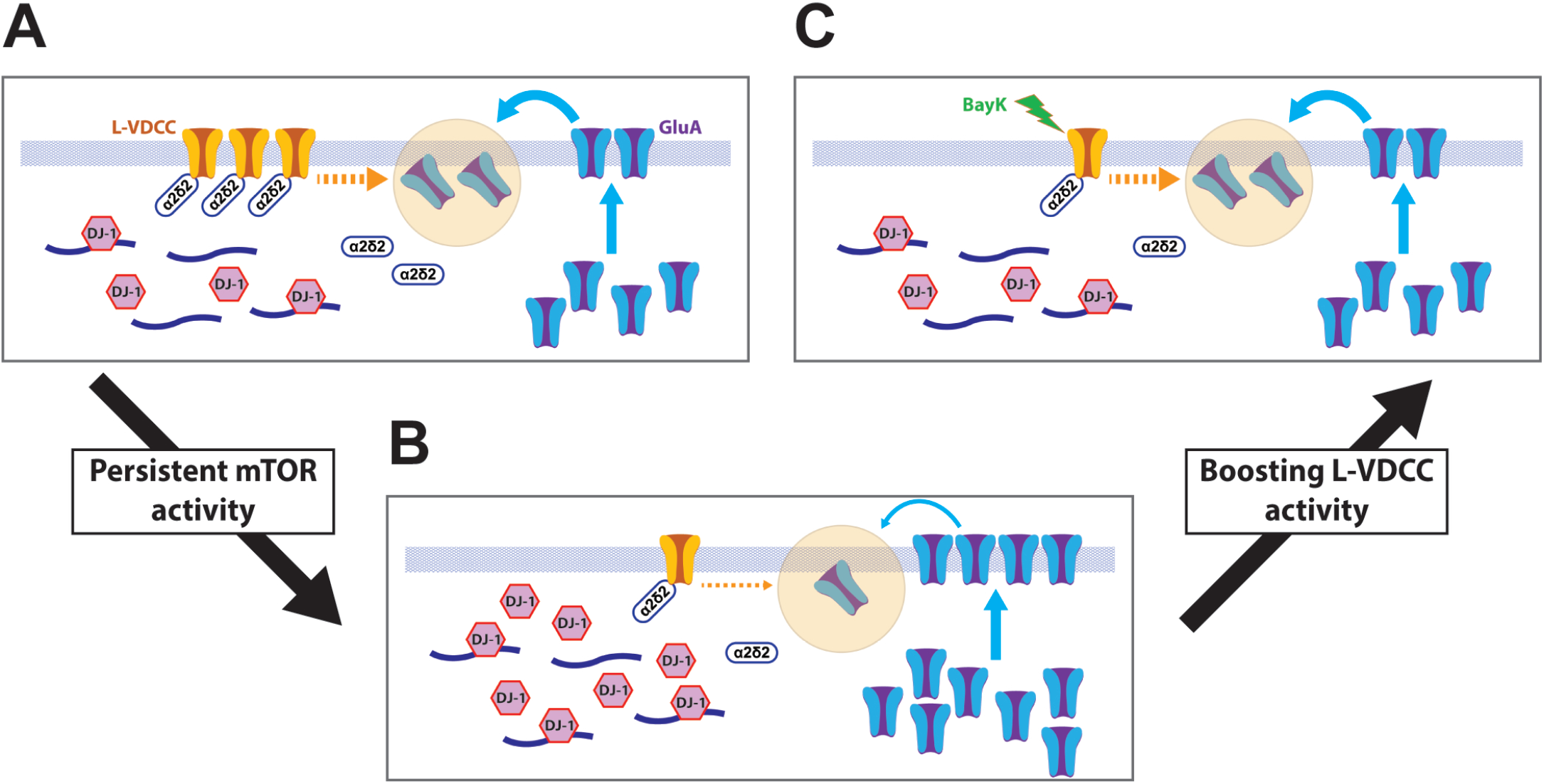
Working model underlying L-VDCC-mediated neuropathophysiology in TS. **(A)** Under normal conditions, the dynamic state of mTOR activity allows DJ-1-regulated events (L-VDCC activity and α2δ2 expression) in dendrites to respond to synaptic stimuli accordingly. L-VDCC activity modulates GluA expression on the dendritic membrane. α2δ2 influences the stability and trafficking of L-VDCC to the membrane surface. **(B)** In a persistent active mTOR state, elevated DJ-1 represses L-VDCC activity and diminishes α2δ2 level. Weakened L-VDCC activity, either through DJ-1/L-VDCC or DJ-1/ α2δ2 interaction or both, promotes more expression of GluA on the membrane. Consequently, overabundance of GluA favors the generation of seizures that can lower life expectancy. **(C)** Boosting L-VDCC activity by BayK in a persistently active mTOR state (e.g. TS) may diminish seizures and improve quality of life potentially through 3 mechanisms: (1) lowering DJ-1 levels (i.e. reversing the effects of excess DJ-1; *see Fig. 1B-C*), (2) decreasing surface expression of GluA (*see Fig. 2B-C*), and (3) reducing mTOR activity (*see Fig. 3D-F*).

Ca^2+^ channels are under-characterized in TS. Our findings that α2δ2 expression is dysregulated in TS and that it regulates GluA expression on the membrane surface, and considering its association with calcium channels argue the value of investigating the calcium channel proteome in TS and other mTORopathies (**Fig. 2**). Furthermore, as BayK does not fully suppress pS6 levels, consideration of L-VDCC activation as a therapeutic avenue should be investigated since this pathway, unlike rapamycin, may still be able to promote mTORC1-dependent syntheses of proteins that may be integral in maintaining normal synaptic function (9, 11). Current therapeutic mitigation of seizures and TS focuses on mTOR and GluA (1, 37, 40-43). Their broad influence and psychiatric side effects, however, limit their clinical utility. Our collective mechanistic and functional findings downstream of mTOR offer another pathway to alleviate TS-associated neurological disorders.

## Methods

### Connectivity Map (CMAP) linked user environment (CLUE) analysis

To identify perturbagens that can elicit upregulation of active-mTORC1-repressed proteins, we seeded CLUE with a set of proteins whose expression levels in the PSD increase with rapamycin, as determined by Niere et al (2016) (9), and whose mRNAs contain putative DJ-1 binding sites (20, 21, 44).

### Generation of TS mice

*Tsc1*^*tm1Djk*^*/J* mice (Jackson Laboratory, Bar Harbor, ME) were used for *in vivo* and *in vitro* experiments. Recombinant adeno-associated virus coding for Cre recombinase (AAV-Cre-GFP) was introduced in *Tsc1*^*tm1Djk*^*/J* mice to knockout the *Tsc1* gene and generate the TS mice. AAV-GFP (vector) was introduced to serve as control (9). A synapsin promoter was used to drive both AAV-GFP and AAV-CRE-GFP.

### Primary neuronal cell culture

Embryonic rat pups (E17–18) were used to generate dissociated hippocampal neurons. Rat neuronal cultures were used for experiments at 18–25 days *in vitro* (DIV). For *Tsc1* cultures, neurons were prepared from male and female postnatal (0–3 days) *Tsc1* conditional knockout pups similar to Niere *et al* (45).

### Pharmacology

(S)-(-)-Bay-K-8644 (Tocris) dissolved in DMSO (Sigma Aldrich) was used for experiments involving BayK. In **Fig. 1 to 3F**, a final concentration of 0.2-2.0 mg/kg of BayK was injected intrapertioneally. These concentrations lower DJ-1 levels in TS mice. As 0.2 and 1.0 mg/kg BayK abates DJ-1 expression in TS, this concentration was used in **Fig. 3G to 4**. Nimodipine (Tocris), dissolved in DMSO, was used at a final concentration of 5 μM.

### Transfection

Primary hippocampal neurons were transiently transfected on DIV 14 with α2δ2 (Addgene) or pcDNA plus RFP plasmid. α2δ2 HA pMT2 was a gift from Annette Dolphin (46). Hippocampal cultures were transfected with Lipofectamine 2000 (Invitrogen) according to the manufacturer’s instructions. Transfection was allowed to occur for 2.5 hours. Cells were processed for immunohistochemistry on DIV18.

### Immunohistochemistry (IHC)

Dissociated hippocampal neurons were fixed in 4% paraformaldehyde for 20 minutes at room temperature and permeabilized in 0.2% Triton X-100 for 10 minutes(45). Fixed cells were incubated overnight in 4°C with the following primary antibodies: rabbit anti-α2δ2 (1:500, Alomone), rabbit anti-DJ-1 (1:1000, Novus), chicken anti-Map2 (1:1000, Aves), mouse anti-Map2 (1:1000, Abcam). For detection of surface GluA1 in rat hippocampal neurons, the cells were fixed in 4% paraformaldehyde for 8 minutes on ice to prevent permeabilization. After blocking in 8% normal goat serum (NGS) for an hour at RT, the cells were incubated with a mouse anti-GluA1 N-terminal antibody (1:100, Millipore) overnight at 4°C. The cells were then washed with PBS, permeabilized for 5 minutes and incubated with either chicken or mouse anti-Map2 for 1 hr at RT. Appropriate secondary antibodies (1:500, Life Technologies) were then used. For detection of surface GluA1 in mouse WT or Tsc1 KO neurons, the cells were incubated with a rabbit anti-GluA1 N-terminal antibody (Calbiochem) for 10 minutes at 37°C. The cells were then fixed on ice for 5 minutes and then incubated with an AlexaFluor647 for 1hr at RT to saturate surface GluA1. Permeabilization and blocking was then performed followed by incubation with chicken anti-Map2 ON at 4°C. Appropriate secondary antibodies (1:500, Life Technologies) were used—AlexaFluor488 (AF488), AF647, AF405—after overnight primary antibody incubation.

### Western blots

#### Synaptoneurosome preparation

Hippocampi were homogenized with lysis buffer composed of Tris-Base with Halt™ Phosphatase and Protease Inhibitor Cocktail (Thermo Scientific). Homogenates were filtered through 100μm cell strainer and 5μm syringe filter, sequentially, and centrifuged for 20 minutes in 4°C at 14,000 xg. The pellet was retained, solubilized in radioimmunoprecipitation (RIPA) buffer, and centrifuged for 10 minutes in 4°C at 14,000 xg. Pierce™ BCA Protein Assay Kit (Thermo Scientific) was used to quantify protein in the collected supernatant.

#### Western blot analysis

SDS-polyacrylamide gel electrophoresis (PAGE) loading buffer using 4x Laemmli Sample Buffer (Bio-Rad) was added to protein samples. 20 to 50 μg protein from each sample were resolved in 10% SDS-polyacrylamide gel and transferred onto 0.2 μm nitrocellulose membranes (Bio-Rad). 5% nonfat dry milk in Tris-buffered saline containing 0.1% Tween 20 was use to block the membranes. To visualize the proteins, we used: mouse anti-DJ1 (1:2000; Novus Biologicals), rabbit anti-phospho-GluA1 (Ser845, 1:1000, R&D Systems), mouse anti-GluA1 (1:2000, Neuromab), mouse anti-ribosomal S6 (1:1000, Cell Signaling), rabbit anti-phospho-S6 (1:1000, Cell Signaling), and mouse anti-tubulin (1:20,000; Abcam). Fluorescence-conjugated secondary antibodies (AF680, Life Technologies; AF800, LiCor; 1:5000) were used to visualize the proteins of interest. Immunoblots were obtained using the Odyssey CLx infrared imaging system. Densitometric quantification of proteins was performed using ImageJ (National Institutes of Health) software.

## Acknowledgments

We would like to thank Dr. Sanjeev Namjoshi for his helpful discussions regarding the connectivity mapping. This study was supported by National Institutes of Health NINDS NS105005 (KRG); the National Science Foundation IOS 1355158 and 1724812 (KRG), Postdoctoral Research Fellowship in Biology DBI-1306528 (FN) and DBI-1103738 (LPC); Alzheimer’s Association AARF-19-614794 (FN); Department of Defense, United States Army Medical Research and Materiel Command USAMRMC Award W81XWH-14-1-0061 and W81XWH-19-1-0202 (KRG). National Institute of Aging Wake Forest School of Medicine Alzheimer’s Disease Research Center Pilot Grant P30AG049638 (KRG).

## Author contributions

FN, LPC, and KRG designed research. FN, LPC, and WCT conducted experiments and analyzed data. FN, LPC, and KRG wrote the manuscript.

## Notes

### Competing Interest Statement

The authors have declared no competing interest.

